# COMPUTATIONAL IDENTIFICATION VALIDATION AND STRUCTURAL CHARACTERIZATION OF SOME POTENTIAL CANDIDATE GENES FOR DIABETES MELLITUS

**DOI:** 10.1101/2022.08.26.505395

**Authors:** Dhananjay Kumar Tanty, Prachi Rani Sahu, Swagatika Behera, Ranjit Mohapatra, Pabitra Mohan Behera, Susanta Kumar Sahu

**Author notes:** Corresponding Authors: Susanta Kumar Sahu and Pabitra Mohan Behera Corresponding author’s address: Dr. Susanta Kumar Sahu, Reader, University Department of Pharmaceutical Sciences, Utkal University, Vani Vihar, Bhubaneswar, Odisha, Pin-751004., Dr. Pabitra Mohan Behera, Application Scientist, Institute of Computational Biology and Bioinformatics, BIOPRUDENCE RESEARCH INNOVATIONS LLP, Bhubaneswar, Odisha, Pin-751002., Contact No. First Authors: Dhananjay Kumar Tanty and Prachi Rani Sahu.

## Abstract

Diabetes mellitus seems to be a complex metabolic disorder due to its association with several complications like cardiovascular, ocular, neurologic, skeletal, hepatic and renal abnormalities etc. Current estimations by WHO suggest that most of the low and middle-income countries of the world are worst affected by this disorder with a prediction that the prevalence may double between 2020 and 2030. Thus both doctors and scientists across the globe are involved in research to disclose the complex genetics of this disorder associated with several environmental and demographic factors. In the last 10 years, several predictions have been made in the lane of omics approaches and computational biology which makes the process quite generous. In the current work, we present a computational analysis of potential candidate genes for diabetes mellitus and their differential expressions in targeted human tissue systems. About 220 reported genes for diabetes mellitus were selected for the study and their protein-protein interaction network (5090 nodes) was extracted using medium-confidence interactions of the HIPPIE database. From the network, the top 10% (509) genes were categorised as hub genes after calculation of about 11 centralities, their consensus ranking and rank correlations. The same set of 220 genes was used for gene ontology enrichment analysis featuring about 1483 genes. About 89 candidate genes were predicted for diabetes mellitus and their differential expressions were studied in adipose, pancreas, skeletal, hepatic and renal tissue systems using the information from NCBI GEOdatabase. Then the differentially expressed gene sets for each tissue system were further validated by fetching them in the potential clusters of the PPI network designed earlier with their functional enrichment analysis using information from the STRINGS database. About 77 genes were prioritized with help of our scoring system and their structural characterization was done with protein centric annotations from UniProtKB database information and molecular model building. We hope our findings are helpful in understanding the expression of diabetes-related genes in different human tissue systems which may lead to the design of newer therapeutic strategies.

## 1. Introduction

The human biological system seems to be a complex of metabolic processes and each such process is suitably regulated by a number of functional genes and corresponding proteins. The application of computational techniques was crucial to understanding the working of such systems which leads the advent of systems biology featuring the study of various interactions of genes, proteins, transcription factors, small molecules, metabolites, and their relationships to complex physiological and pathological processes. Network biology seems to be a primary method used for studying numerous interactions which occur inside the cell and their role in various complex biological processes [1]. It also reveals the relationships between different biological components and tries to fetch the existence of any physical or functional relationships with other components to be identified or interrogated [2, 3]. Besides these it also provides information related to the pathobiology of disorders, describing the pathways associated with a particular disease process and prioritization of candidate genes causing a particular disease. These credentials of network biology have been exploited in the generation of so-called disease networks featuring a list of disease-gene associations explaining the relationship between genes and corresponding genetic disorders [4]. The human diseasome is designed to reflect the cohesive relationship between the disease genes and their products. It also signifies the increased propensity of disease genes towards their products and the tendency to express in specific tissues with common cellular and functional characteristics [5]. Biological networks focusing on a particular disease are now vastly used due to their predictive power in describing the protein-protein interactions, physical interactions, the impact of biological markers, and the pathophysiology of the concerned disease. Some examples are the implementation of gene networks in different types of tumors like colorectal [6], ovarian [7], and breast cancers [8]. Other such applications are in neurological disorders like Parkinson’s [9] and Alzheimer’s disease [10], and neuropsychiatric disorders like autoimmune disease [11] and hematologic disease [12]. The overwhelming increase in several lifestyle disorders like coronary heart disease [13], fatty liver diseases [14], and endocrine/metabolic disorders demands the application of network biology in understanding their pathogenesis and disease-gene interactions. Diabetes seems to be one of the major human endocrine disorders, which may onset during infancy, passes through childhood and exists till late adulthood. In the recent past prevalence and significance of different diabetes forms in these three main stages of human life have been studied thoroughly with the use of network analysis. Congenital hyperinsulinism (CHI) is a metabolic disorder that appears during the infancy stage of humans characterized by dysregulation of excess insulin secretion from pancreatic beta-cells and leads to critical hypoglycemia [15]. The computational biological study assisted by network biology revealed about nine genes to be involved in CHI which can be used as biomarkers for diagnosis and management purposes [16, 17]. The application of network analysis also revealed that human growth is linked to the changes in the expression of a certain set of genes within evolutionarily conserved growth pathways [18].

Being inspired by the application of systems biology in different diseases as mentioned above we present computational identification validation and structural characterization of candidate genes for diabetes mellitus by integrating the validated information from various public databases. The prediction of candidate genes was based on the assumption that one particular gene is not potent enough for the causal of a disease, rather genes or groups of genes are responsible for the cause. Again the genes that are in the close proximity regions of a particular disease-causing gene may be involved in the onset of that disease, thus hypothesized as candidate genes.

## 2. Materials and Methods

### 2.1 Materials

#### 2.1.1 Selection of genes associated with diabetes mellitus

The latest version of DisGeNET (v7.0) featuring about 1,134,942 gene-disease associations (GDAs) was searched for the selection of genes associated with diabetes mellitus. The GDAs were reported among 21,671 genes and 30,170 diseases with data reported from expert-curated repositories, GWAS catalogs, animal models, and published literature which made the process more exclusive and reliable [19-21] About 220 such genes were programmatically extracted from the ALL_gene-disease association with disease-gene association score (DisGeNet score) which suggests the confidence score of association of a particular disease with a specific gene.

#### 2.1.2 Selection of human protein-protein interaction network

The latest version of HIPPIE (Human Integrated Protein-Protein Interaction rEference) v2.2 was downloaded featuring about 270000 confidence scored and functionally annotated interactions [22, 23]. The selection of such interactions was crucial because HIPPIE seems to be an important resource for the generation and interpretation of protein-protein interaction networks citing a specific research interest.

### 2.2 Methods

#### 2.2.1 Generation of diabetes genes protein-protein interaction network

All 220 diabetes mellitus-associated genes were used as input for extraction of a diabetes genes protein-protein interaction network. A medium confidence network was extracted with a confidence score(s=0.63) and a layer of (l=1) from the global human protein-protein interaction network selected for the HIPPIE database. The medium confidence was selected to include some reasonable number of interactions rather than most exclusive interactions with high confidence of (s=0.72) and most inclusive interaction with low confidence of (s=0.52). Again the fixing of interaction layer size (l=1) generates a potential network featuring all the interactions achieved by the members of the input set. The concerned network is imported in Cytoscape [24] with a preferred prefuse force directed layout for visualization. About 5090 nodes and 10368 edges were observed in the network form in which the largest subnetwork was extracted with 5083 nodes and 10362 edges.

#### 2.2.2 Gene Ontology Enrichment analysis of the diabetic genes

The gene ontology enrichment analysis requires two sets of genes (target set and background set) as input and the enrichment is fetched in the target set in comparison with the background set. The Gorilla (GeneOntology enRIchment anaLysis and visuaLizAtion) [25] web-based application was used for the purpose with all 220 diabetes genes as the target set and all 5090 nodes of the diabetes genes PPI network as the background set. All three ontology terms (process, function, and component) were selected for fetching enriched GO terms with a p-value threshold of 10-3. The output was customized to be saved in Excel, the inclusion of both unresolved and duplicate genes and direct opening of resulting GO terms with their p-value in the REVIGO (REduce VIsualize Gene Ontology) tool [26]. After selection of significantly enriched GO terms for all three ontology terms and removal of redundancy the genes associated with the GO terms were fetched in file goa_human.gaf downloaded from Gene Ontology Resource [27, 28].

#### 2.2.3 Centrality calculation of diabetes genes PPI network and selection of hub genes

The calculation of centralities of constituent nodes of a network is important as it measures and assigns the ranking of the concerned nodes corresponding to their network positions. Applications of centrality include the identification of the most important or influential nodes (hubs) in the network. The medium confidence diabetes genes PPI network was imported in Cytoscape with default layout and about eleven different centralities were calculated with the cytoHubba app [29]. These eleven centralities were based on either the local-based method (Degree) or the global-based method (shortest path). The local-based centralities were Deg (Degree), MNC (Maximum Neighborhood Component), DMNC (Density of Maximum Neighborhood Component), and MCC (MaximalClique Centrality). The global-based centralities were Clo (Closeness), EC (EcCentricity), Rad (Radiality), BN (BottleNeck), Str(Stress), BC (Betweenness), and EPC (Edge Percolated Component). Out of these eleven centralities, MCC seems to be a better performer than the others with a capability to capture more essential nodes or hubs in the top-ranked lists in both high degree and low degree of genes. The ranked list of all 5090 nodes by eleven different centralities was downloaded and analyzed for the selection of hub genes. It was observed that all 5090 nodes have different rankings for eleven different centralities. Thus a consensus-based ranking was targeted in which all eleven different rankings were sorted by the nodes and for each node, all ranks were added to fetch the consensus ranking score. Then the consensus ranking was sorted by ascending order and nodes achieving the top ten percent of the consensus ranking were designated as hub genes.

#### 2.2.4 Prediction of candidate genes for diabetes mellitus

The prediction of potential candidate genes for diabetes mellitus was achieved by defining three sets of genes. The diabetes mellitus-related genes selected from the DisGeNET database were designated as set1. The genes that were associated with significantly enriched GO terms after gene ontology enrichment analysis of diabetes mellitus genes defined in set1 were designated as set2. The prediction of hub genes after centrality calculations and consensus ranking were designated as set3. The candidate genes were defined as the genes that were not in set1 but common between set2 and set3. In elaboration, the genes that were not categorized as diabetes mellitus genes but are highly enriched with the gene ontology terms defined for diabetes mellitus and designated as hub genes in the diabetes gene PPI network were the candidate genes.

#### 2.2.5 Differential expression analysis of candidate genes in selective tissue systems

The NCBI GEO (Gene Expression Omnibus) [30, 31] database was searched with the keyword “diabetes mellitus” from which about 59 human-specific datasets were selected. All these 59 datasets were analyzed for the selection of tissue-specific GEO series. Thus the expression series selected for five tissue systems were adipose (GSE16415, GSE29226, GSE40234), pancreas (GSE20966, GSE25724, GSE38642), skeletal (GSE29221), hepatic (GSE1009, GSE3308) and renal (GSE15653, GSE23343). Each GEO series file was analyzed with an interactive web tool GEO2R (https://www.ncbi.nlm.nih.gov/geo/geo2r) for comparison of two groups (diabetic and normal) of samples defined in the respective GEO series for identification of genes that are differentially expressed across different experimental conditions. The analysis is further customized with the selection of Benjamini & Hochberg false discovery rate, auto-detection for log transformation of values, without application of limma precision weights (vooma), no force normalization of the expression data, and with default adj-P-value significance level cut-off of 0.05. The concerned R script was modified for the selection of the top 10000 genes and the output was exported in excel format for further analysis.

#### 2.2.6 Fetching of candidate genes in potential clusters of diabetes PPI network

Prediction of potential clusters in the complex PPI networks is crucial in the identification of groups of similar entities (genes/proteins) with similar structures and functions. The clustering algorithms designed for the purpose mostly rely on mathematical and logical approximations to build meaningful relationships between the entities under consideration. The MCODE (Molecular Complex Detection) [32] application of Cytoscape was used for the detection of densely connected regions in the diabetes PPI network as clusters. The clusters were predicted by customizing the network scoring parameters and cluster finding parameters in two iterations. In the first iteration the clusters were fetched with network scoring parameters as the inclusion of loops and degree cutoff value of two and with cluster finding parameters as Haircut with Node Score Cutoff value of 0.2, K-core value of 2, and Max. Depth of 100. The second iteration was run with similar parameters as selected for the first iteration with the inclusion of Fluff and Node Density Cutoff of 0.1.

### 2.2.7 Functional enrichment analysis of candidate genes

The functional enrichment of 89 candidate genes was done by querying their names in the STRING protein query box of the STRING app [33]. The full STRING network was retrieved with a confidence score cut-off value of 0.4 and using smart delimiters. The functional enrichment data were retrieved by selecting all candidate genes and the corresponding STRING network as background. The enrichment data for biological process, molecular function, tissue, and KEGG pathway were exported for further analysis. The redundant terms were removed with a redundancy cutoff of 0.5 prior to exportation as individual files.

#### 2.2.8 Prediction of potential mutations in the candidate genes

Mutations in the genes are the start point of several complications as they suitably modulate their regular expressions. They are either benign with no effect on the genetic expression or deleterious with extreme effects on the expression thus triggering structural and functional modifications. All 89 candidate genes were searched in the UniProtKB [34] for the selection of their reported natural variants. The reported natural variants were further searched for their association with disease-related mutations in the PredictSNP server [35]. The prediction of amino acid substitutions in the concerned gene was searched by inserting the corresponding protein sequence in FASTA format and defining the mutation positions characterized by their natural variants. All seven tools were selected for evaluation of the mutations.

#### 2.2.9 Structural characterization of candidate genes

The structural characterization was done by designing computational models of the deleterious natural variants of genes. In most cases, the models were developed by homology modeling strategy which included searching for suitable templates, alignment of the target and template, model building by Modeller 10.2 [36-39] and evaluation by predicting Ramachandran plots through PROCHECK [40]. The genes which do not have a suitable template for model building their secondary structure were predicted using the utilities of I-TASSER (Iterative Threading ASSEmbly Refinement) threading server [41-43].

The whole methodologies followed in the current work are summarized as a flow-chart mentioned in the figure-1.

**Figure-1:**
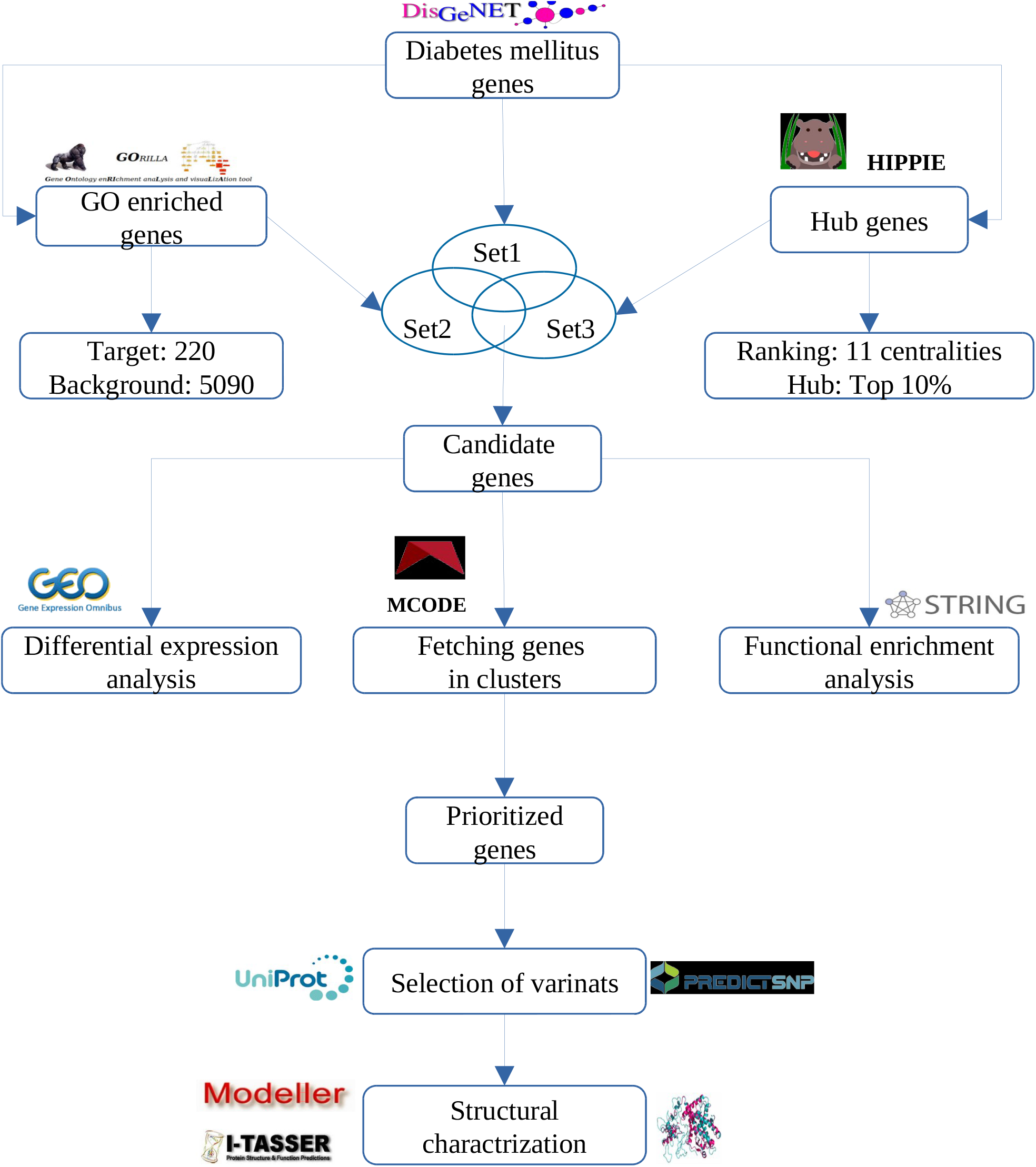
The flow-chart showing methods followed in the current research work

## 3. Result and discussions

### 3.1 Prediction of potential candidate genes for diabetes mellitus

The medium confidence diabetes mellitus network extracted from the protein-protein interactions of the HIPPIE database was analyzed with the analysis network tool Cytoscape. The important topological parameters associated with the network were network diameter of 9, network radius of 5, the clustering coefficient of 0.060, network density of 0.001, network heterogeneity of 5.968, and network centralization of 0.258. The degrees associated with 5090 nodes were selected for node degree distribution by calculating the frequency of each degree and the fraction of nodes of the network associated with it. The graph shown in Fig-2A represents node degree as K and fraction of nodes as p(K). The exponent value of 1.147 in the equation indicates that the network is a scale-free and robust biological network.

**Figure-2:**
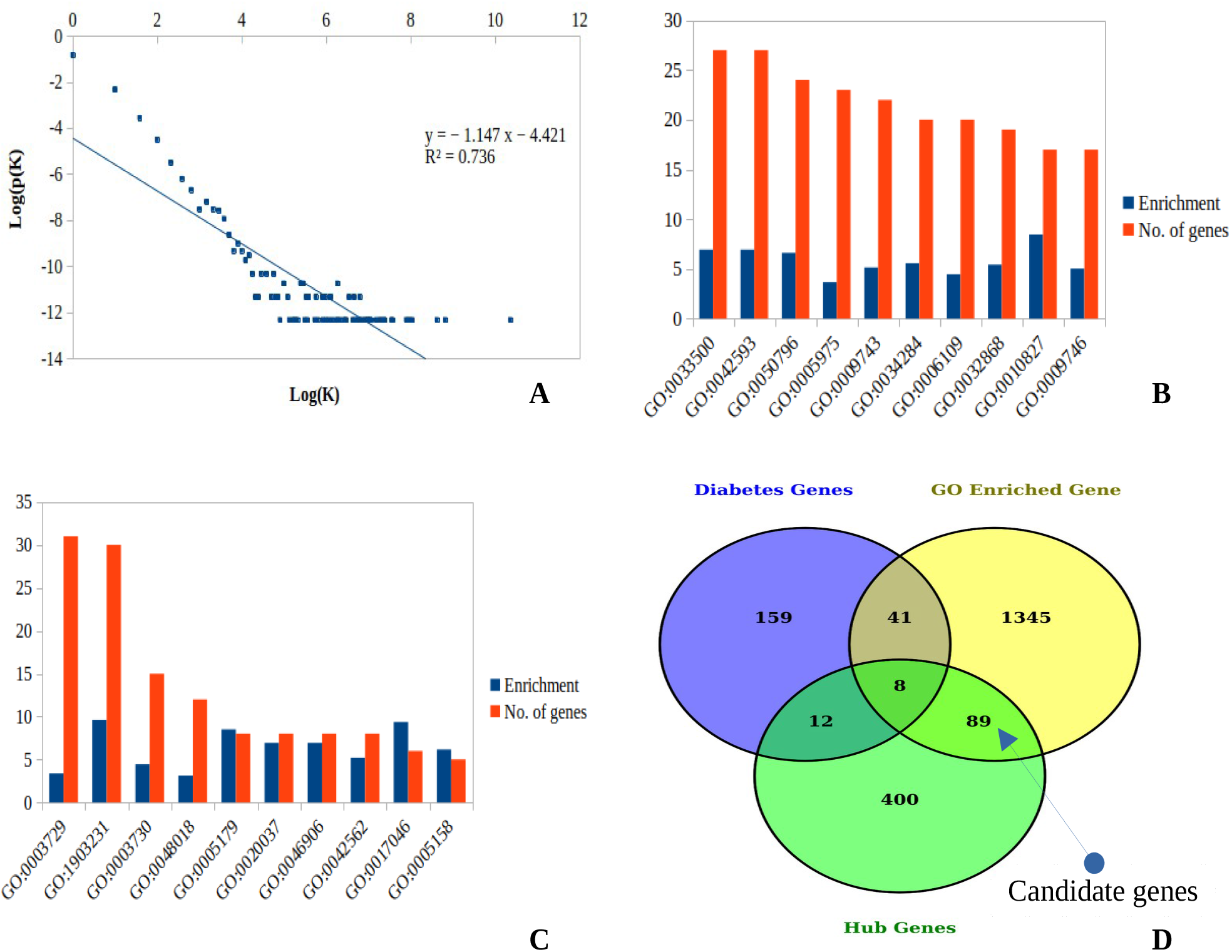
**A:** The graph of node degree distribution with degree in X-axes (Log(K)) and fraction of nodes in the network in the Y-axes (Log(p(K))), **B:** The GO terms enriched for biological process with enrichment and no. of genes, **C:** The GO terms enriched for molecular function with enrichment and no. of genes, **D:** The venn diagram showing the 89 candiadte genes for diabetes mellitus.

The gene ontology enrichment analysis generated about 435 enriched GO terms sorted by P-value, FDR q-value, and enrichment. More precisely there were 419, 11, and 5 GO terms enriched for biological process, molecular function, and cellular component respectively. The identification of enriched GO terms is based on the hypergeometric distribution of total N genes from which B is associated with a particular GO term (background set) and from a total of n genes in which the probability of b or more genes are associated with that particular GO term (target set). Thus a cut of the threshold of (49 ≤ b ≥4) was used for the selection of significant GO terms which generated about 362 GO terms of which 349, 10, and 3 GO terms were enriched for biological processes, molecular function, and cellular component respectively. The top ten significant GO terms enriched for biological process and molecular function are shown in Fig-2B and Fig-2C. Only 182 GO terms were left after removal of redundant GO terms which were further searched in the file goa_human.gaf for the association of genes that produced about 1483 unique genes.

The consensus-based ranking of the global diabetes protein-protein network ranked the whole 5090 nodes by eleven different centralities. This strategy provided a better observation of nodes as hubs as they were ranked by consensus ranking (ranking of ranks) rather than individual ranking by a particular centrality. This top ten percent of nodes (509) with consensus ranking were selected as hub genes.

The potential candidate genes for diabetes mellitus were predicted from three sets of genes such as 220 diabetes mellitus-associated genes selected from the DisGeNET database (Set1), 1483 GO enriched genes as reported from gene ontology enrichment analysis (Set2), and 509 hub genes of medium confidence diabetes network (Set3). About 89 genes were predicted as potential candidate genes for diabetes mellitus, those were previously not reported as diabetic genes but are highly GO enriched genes and hub genes. (Fig-2D)

### 3.2 Differential expression of candidate genes in selected tissue systems

Prior to analysis, the concerned r scripts from the GEO2R tool were modified with the inclusion of different libraries like GEOquery, limma, umap, dplyr, pheatmap, readr, readxl, and EnhancedVolcano, and top 10000 genes were exported as output with adjusted_P_value, P_value, and logFC scores. All 89 candidate genes were searched in the resulting excel files and their scores achieved in each GEO series are tabulated (adjusted_P_val>0.05 and logFC>1) in form of a heat-map shown in Fig-3A. The genes with logFC>1 were considered up-regulated and logFC<1 were considered as down-regulated in particular tissue systems. About 57 candidate genes were associated with the significant values of logFC thus appeared in the heat map but their scores were not significant for the GEO series GSE12643 and GSE25462 of the skeletal tissue system. The most differentially expressed genes in different tissue systems were adipose (22 genes), hepatic (4 genes), pancreatic (34 genes), renal (26 genes), and skeletal (7 genes) Fig-3B.

**Figure-3:**
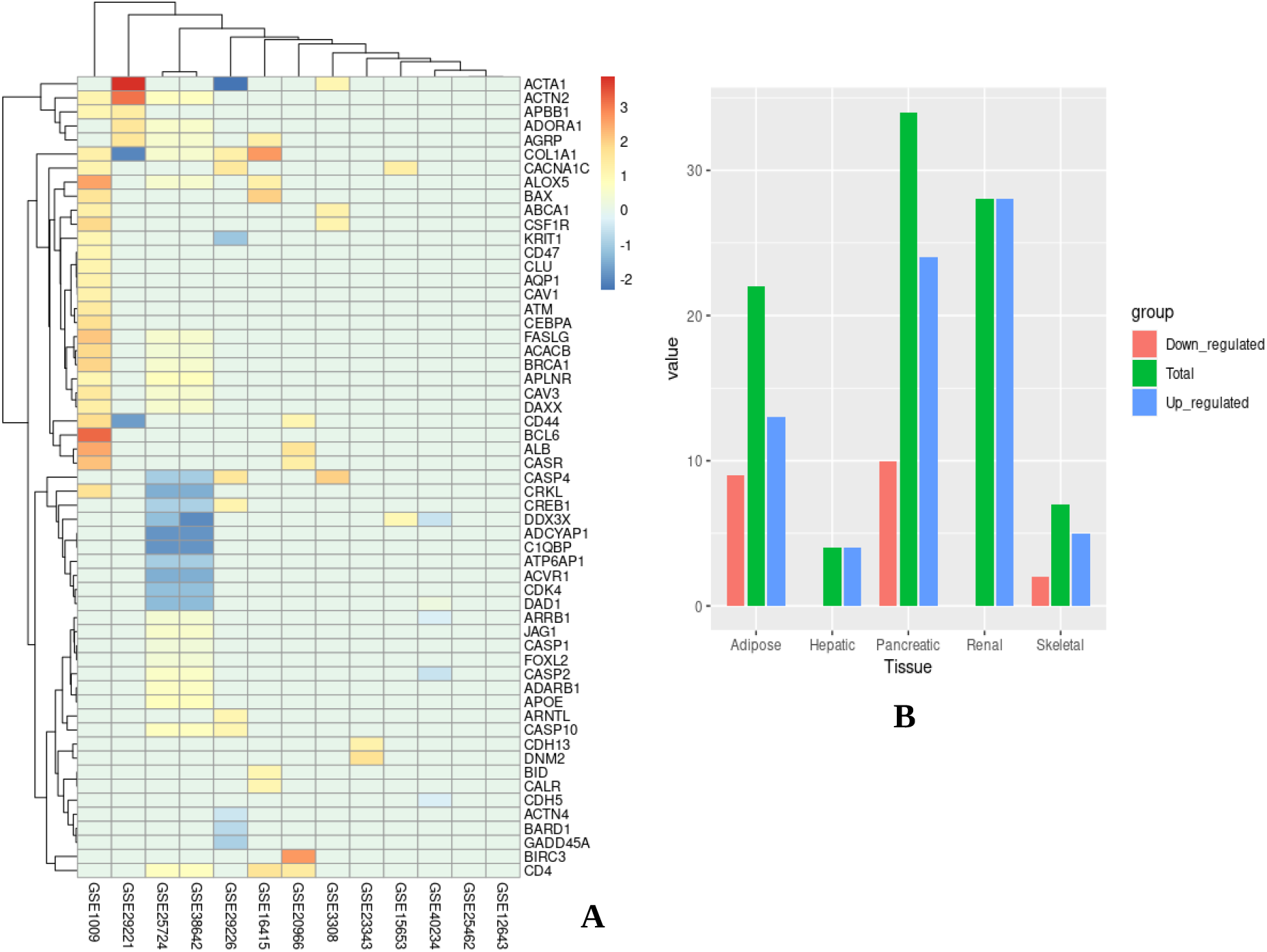
**A:** The heat map shows expression of 57 candidate genes based on their logFC values, **B:** The bar plot shows the number of up-regulated, down-regulated and total candidate genes in five different tissue systems.

### 3.3 Prediction of candidate genes in the clusters

About five different clusters were generated for each iteration of clustering, ranked by their clustering score. The top two clusters from each iteration were selected for fetching of both candidate genes, diabetes genes, and the precision of finding as shown in Table-1. The occurrence of less number of nodes in first iteration clusters (haircut clusters 1 and 2) is characterized by the removal of nodes with single connections to the cluster, although they satisfy the degree cutoff parameter during clustering and scoring. The occurrence of more nodes in the case of the second iteration clusters (haircut fluff clusters 1 and 2) is based on the fact that when fluff is turned on the MCODE program has the freedom of expanding the cluster core by one neighbour shell outwards (layer-1) with specified node density cutoff.

**Table-1:** The top two clusters of each iteration with their topological features

| Sl. No. | Cluster type | No. of nodes | No. of edges | Candidate genes | Diabetes genes | Precision |
| --- | --- | --- | --- | --- | --- | --- |
| 1 | Haircut cluster 1 | 31 | 56 | 3 | 11 | 0.27 |
| 2 | Haircut cluster 2 | 27 | 45 | 1 | 6 | 0.16 |
| 3 | Haircut fluff cluster1 | 104 | 216 | 1 | 8 | 0.11 |
| 4 | Haircut fluff cluster2 | 691 | 843 | 15 | 6 | 2.62 |

The network described by the first cluster was observed to have about 31 nodes and 56 edges with a score of 3.733. About 3 candidate genes (AKT1, BID, and CREBBP) and 11 diabetes genes (AKT2, BCL2, CCND2, CASP3, FAS, HBA1, HMGA1, HNF4A, PPARG, PPARGC1A, and PROX1) were mapped with a precision of 0.27. In the network both AKT1 and BID are associated with a degree of three and CREBBP is associated with a degree of four (Fig-3A). The network described by the second cluster was observed to have 27 nodes and 45 edges with a score of 3.462. Only 1 candidate gene (ARRB2) and 6 (BRAF, EDNRA, EGFR, IRS2, HK1, and MRAS) diabetes genes were mapped with a precision of 0.16. The diabetes genes like BRAF, EGFR, and HK1 occupied the crucial positions with the most degrees in comparison to other members, and the single candidate gene was associated with a degree of 2 (Fig-3B). The network described by the third cluster was observed to have 104 nodes and 216 edges with a score of 4.114. Only 1 candidate gene (ADARB1) and 8 (AP3S2, CMIP, DGKD, MIR98, MIR27A, MIR205, MIR221, and ZFAND3) diabetes genes were mapped with a precision of 0.11. The network described by the fourth cluster was observed to have about 691 nodes and 843 edges with a score of 2.436. About 15 candidate genes (ABCA1, ACTG1, ACTA1, ALB, ALOX5, ATF4, BARD1, BRCA1, C1QBP, CDK4, CLU, CREBBP, CTNNB1, DAXX, and DHX9) and 6 diabetes genes (GCKR, NEUROD1, NFKB1, PROX1, RELA, and SLC1A2) were mapped with a precision of 2.62. Topological observation of this cluster suggests the occurrence of three major sub-clusters of which two have a central diabetes gene and other associated candidate genes (Fig-3C). Again a group of four diabetes genes like GCKR, NEUROD1, PROX1, and SLC1A2 lie in the vicinity joining two major sub-clusters. These are supposed to maintain the integrity of the cluster along with the other genes connecting the major sub-clusters.

### 3.4 Functional enrichment of candidate genes

All 89 candidate genes were mapped by the STRING database which produced a protein-protein network of 89 nodes and 466 edges. The network was characterized by three singleton nodes (ACACB, ADARB1, and ATP6AP1) with an average node degree of 10.5, the average local clustering coefficient of 0.524, and PPI enrichment of (p-value < 1.0e-16) (Fig-5A). There were more interactions (466) between the queried proteins than expected (202) which shows that the proteins are partially connected as a group to form a biological network. The whole enrichment data was exported as a tabular format which contained about 731 rows of information after the removal of redundant terms. The number of GO terms enriched for biological process (449), molecular function (39), and cellular component (51) were selected for mapping of the number of enriched genes in the PPI network. The top ten GO terms enriched for biological process, molecular function, and cellular component are shown with the number of associated candidate genes in Fig-5B and with transferred FDR values in Fig-5C.

**Figure-4:**
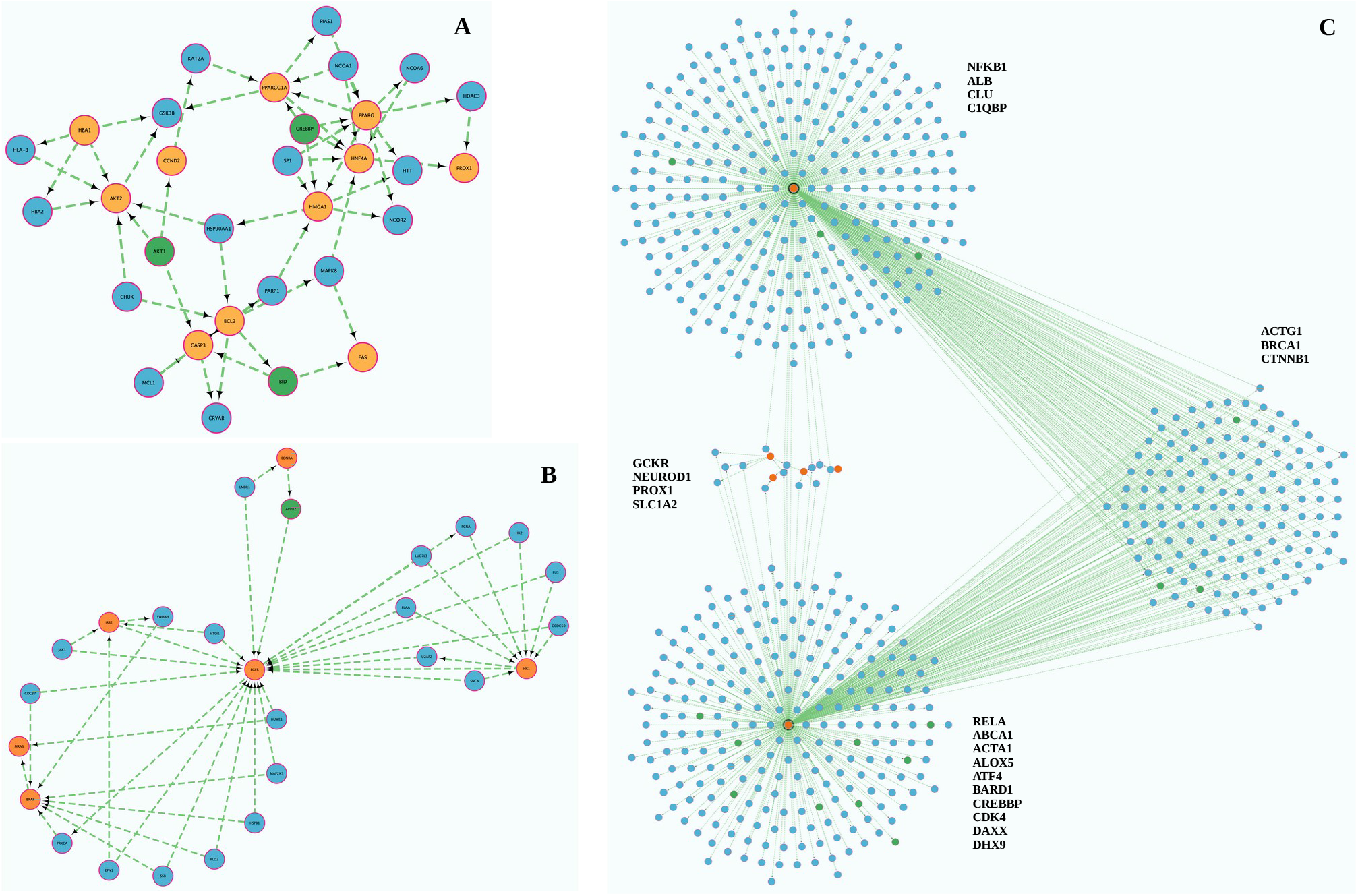
**A:** The haircut cluster1 with 3 candidate genes in green and 11 diabetes genes in crimson colour, **B:** The haircut cluster2 with 1 candidate gene in green and 6 diabetes genes in crimson colour **C:** The haircut fluff cluster2 with 15 candidate genes distributed in different subclusters coloured in green and 6 diabetes genes coloured in crimson.

**Figure-5:**
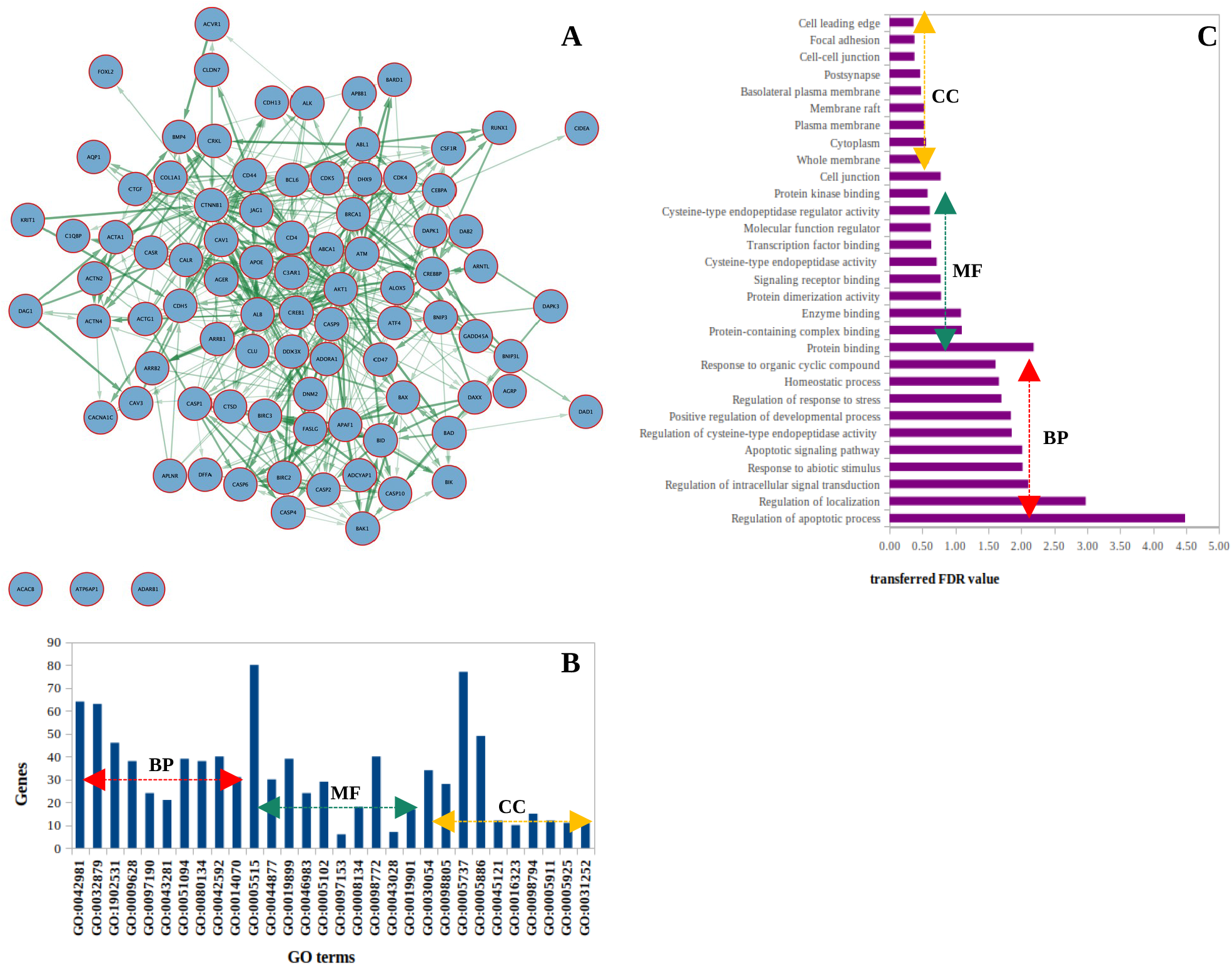
**A:** The PPI network of 89 candidate genes with yFiles organic layout. **B:** The top ten enriched GO terms with number of candidate genes for biological process (BP), molecular function (MF) and cellular component (CC) **C:** The top ten enriched GO terms with tranferred fdr values for biological process (BP), molecular function (MF) and cellular component (CC). Some important GO terms especially enriched for diabetes mellitus were biological processes (glucose homeostasis, response to oxygen-glucose deprivation, and response to glucose), molecular function (protein kinase binding and protein kinase activity), and cellular component (endocrine vesicle). Further analysis of the enrichment result revealed that most of the candidate genes were enriched for five different tissue systems selected for their differential expression analysis described earlier. The number of candidate genes enriched was adipose (17 genes), pancreas (10 genes), skeletal (21 genes), hepatic (35 genes), and renal (15 genes).

The process of identification of 89 candidate genes and their validation by three methods was summarized in the form of a scoring system in which a particular gene is given a score (1) if it follows a method of validation and 0 for not following that particular validation. Then the scores achieved by each candidate gene for three different validation methods were added for the generation of total score. The total score was sorted in descending which categorized the candidate genes into four different groups highly susceptible group (total score=3), moderately susceptible group (total score=2), less susceptible group (total score=1) and false-positive group (total score=0) as shown in Table-2.

**Table-2:** The scores achieved by candidate genes in three different validation processes with a cumulative total score

| Sl. No. | Gene Name | Differential expression | Occurence in cluster | Functional annotation | Total score |
| --- | --- | --- | --- | --- | --- |
| 1 | ABCA1 | 1 | 1 | 1 | 3 |
| 2 | ACTA1 | 1 | 1 | 1 | 3 |
| 3 | ALB | 1 | 1 | 1 | 3 |
| 4 | C1QBP | 1 | 1 | 1 | 3 |
| 5 | CDK4 | 1 | 1 | 1 | 3 |
| 6 | CLU | 1 | 1 | 1 | 3 |
| 7 | ACACB | 1 | 0 | 1 | 2 |
| 8 | ACTG1 | 0 | 1 | 1 | 2 |
| 9 | ACTN2 | 1 | 0 | 1 | 2 |
| 10 | ACTN4 | 1 | 0 | 1 | 2 |
| 11 | ADARB1 | 1 | 1 | 0 | 2 |
| 12 | AKT1 | 0 | 1 | 1 | 2 |
| 13 | ALOX5 | 1 | 1 | 0 | 2 |
| 14 | APOE | 1 | 0 | 1 | 2 |
| 15 | AQP1 | 1 | 0 | 1 | 2 |
| 16 | ARRB2 | 0 | 1 | 1 | 2 |
| 17 | ATF4 | 0 | 1 | 1 | 2 |
| 18 | ATM | 1 | 0 | 1 | 2 |
| 19 | ATP6AP1 | 1 | 0 | 1 | 2 |
| 20 | BARD1 | 1 | 1 | 0 | 2 |
| 21 | BCL6 | 1 | 0 | 1 | 2 |
| 22 | BID | 1 | 1 | 0 | 2 |
| 23 | BIRC3 | 1 | 0 | 1 | 2 |
| 24 | BRCA1 | 1 | 1 | 0 | 2 |
| 25 | CACNA1C | 1 | 0 | 1 | 2 |
| 26 | CALR | 1 | 0 | 1 | 2 |
| 27 | CASR | 1 | 0 | 1 | 2 |
| 28 | CAV1 | 1 | 0 | 1 | 2 |
| 29 | CAV3 | 1 | 0 | 1 | 2 |
| 30 | CD4 | 1 | 0 | 1 | 2 |
| 31 | CD44 | 1 | 0 | 1 | 2 |
| 32 | CD47 | 1 | 0 | 1 | 2 |
| 33 | CDH13 | 1 | 0 | 1 | 2 |
| 34 | CDH5 | 1 | 0 | 1 | 2 |
| 35 | CEBPA | 1 | 0 | 1 | 2 |
| 36 | COL1A1 | 1 | 0 | 1 | 2 |
| 37 | CREB1 | 1 | 0 | 1 | 2 |
| 38 | CREBBP | 0 | 1 | 1 | 2 |
| 39 | CSF1R | 1 | 0 | 1 | 2 |
| 40 | CTNNB1 | 0 | 1 | 1 | 2 |
| 41 | DAXX | 1 | 1 | 0 | 2 |
| 42 | DDX3X | 1 | 0 | 1 | 2 |
| 43 | DHX9 | 0 | 1 | 1 | 2 |
| 44 | GADD45A | 1 | 0 | 1 | 2 |
| 45 | JAG1 | 1 | 0 | 1 | 2 |
| 46 | ABL1 | 0 | 0 | 1 | 1 |
| 47 | ACVR1 | 1 | 0 | 0 | 1 |
| 48 | ADCYAP1 | 1 | 0 | 0 | 1 |
| 49 | ADORA1 | 1 | 0 | 0 | 1 |
| 50 | AGER | 0 | 0 | 1 | 1 |
| 51 | AGRP | 1 | 0 | 0 | 1 |
| 52 | APAF1 | 0 | 0 | 1 | 1 |
| 53 | APBB1 | 1 | 0 | 0 | 1 |
| 54 | APLNR | 1 | 0 | 0 | 1 |
| 55 | ARNTL | 1 | 0 | 0 | 1 |
| 56 | ARRB1 | 1 | 0 | 0 | 1 |
| 57 | BAD | 0 | 0 | 1 | 1 |
| 58 | BAX | 1 | 0 | 0 | 1 |
| 59 | BIRC2 | 0 | 0 | 1 | 1 |
| 60 | BNIP3 | 0 | 0 | 1 | 1 |
| 61 | BNIP3L | 0 | 0 | 1 | 1 |
| 62 | CASP1 | 1 | 0 | 0 | 1 |
| 63 | CASP10 | 1 | 0 | 0 | 1 |
| 64 | CASP2 | 1 | 0 | 0 | 1 |
| 65 | CASP4 | 1 | 0 | 0 | 1 |
| 66 | CASP9 | 0 | 0 | 1 | 1 |
| 67 | CRKL | 1 | 0 | 0 | 1 |
| 68 | CTSD | 0 | 0 | 1 | 1 |
| 69 | DAB2 | 0 | 0 | 1 | 1 |
| 70 | DAD1 | 1 | 0 | 0 | 1 |
| 71 | DAG1 | 0 | 0 | 1 | 1 |
| 72 | DAPK3 | 0 | 0 | 1 | 1 |
| 73 | DFFA | 0 | 0 | 1 | 1 |
| 74 | DNM2 | 1 | 0 | 0 | 1 |
| 75 | FASLG | 1 | 0 | 0 | 1 |
| 76 | FOXL2 | 1 | 0 | 0 | 1 |
| 77 | KRIT1 | 1 | 0 | 0 | 1 |
| 78 | ALK | 0 | 0 | 0 | 0 |
| 79 | BAK1 | 0 | 0 | 0 | 0 |
| 80 | BIK | 0 | 0 | 0 | 0 |
| 81 | BMP4 | 0 | 0 | 0 | 0 |
| 82 | C3AR1 | 0 | 0 | 0 | 0 |
| 83 | CASP6 | 0 | 0 | 0 | 0 |
| 84 | CCN2 | 0 | 0 | 0 | 0 |
| 85 | CDK5 | 0 | 0 | 0 | 0 |
| 86 | CIDEA | 0 | 0 | 0 | 0 |
| 87 | CLDN7 | 0 | 0 | 0 | 0 |
| 88 | DAPK1 | 0 | 0 | 0 | 0 |
| 89 | RUNX1 | 0 | 0 | 0 | 0 |

This short scoring system fine-tuned our analysis to a greater extent by grouping and prioritizing the candidate genes for further structural characterization process. The candidate gene’s protein-protein interaction network generated by the STRING app is then decomposed into four different networks containing the genes described in four groups. The highly susceptible group network had about six candidate genes with one singleton gene (Fig-6B). The moderately susceptible group network was about thirty-nine candidate genes with three singleton genes (Fig-6C). The less susceptible group network had about thirty-two candidate genes with nine singleton genes (Fig-6D). The singleton genes for three groups were retained as such for analysis instead of removing them through a selection of the most significant connected component. The networks defined by high, moderate, and less susceptible groups were further extended by the CyTargetLinker app [44, 45] for fetching associated pathways with the latest available related linksets from the application site. About 55, 303, and 120 different pathways were associated with high, moderate, and less susceptible groups network (Supplementary file-1).

**Figure-6:**
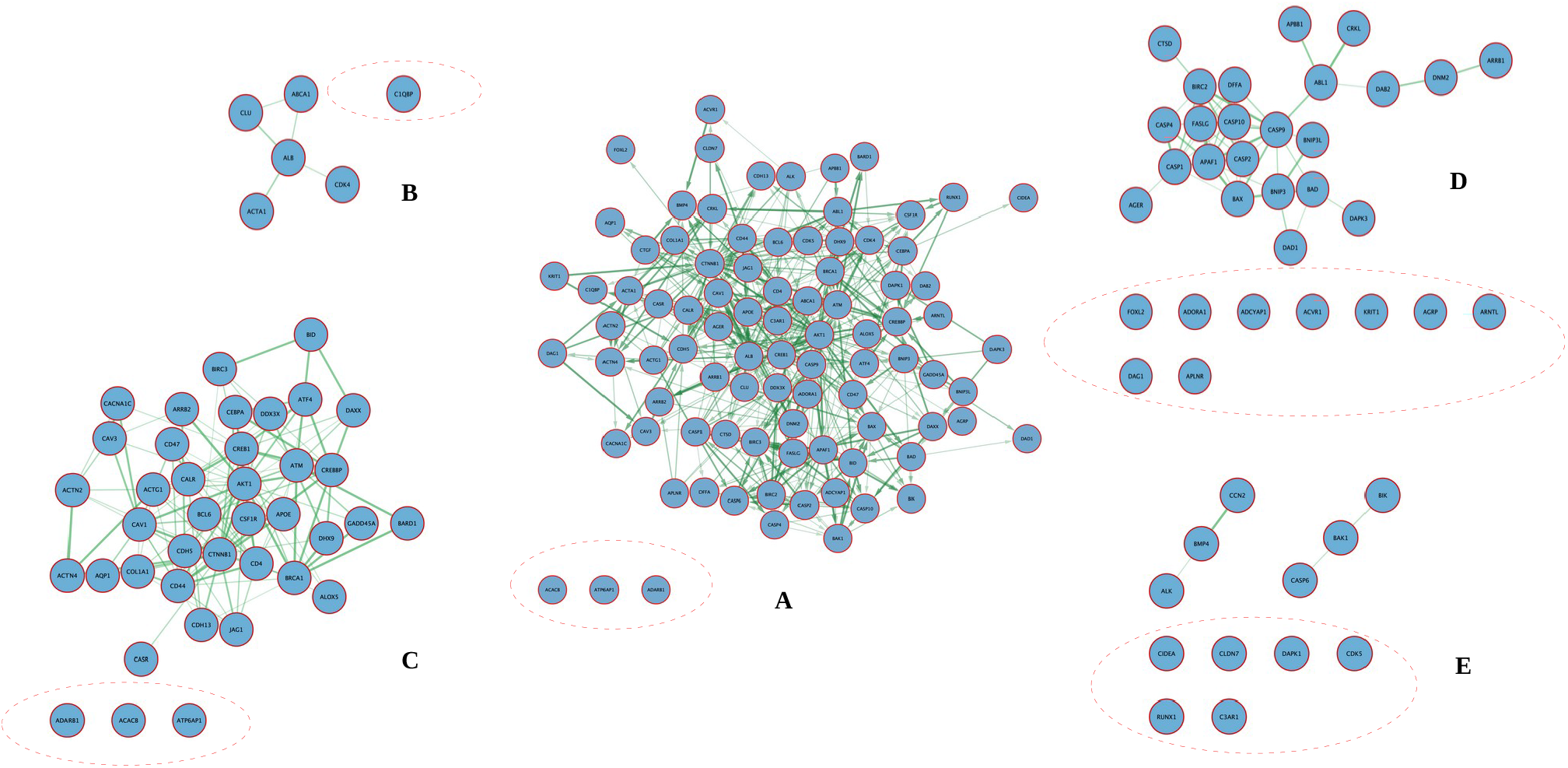
**A:** The PPI network of 89 candidate genes and other four derived groups with singleton genes highlighted in red dotted lines **B:** The highly susceptible genes network **C:** The moderately susceptible genes **D:** The less susceptible genes network **E:** The false positive genes network.

Further in-depth analysis revealed that the highly susceptible gene group were associated with the PPAR-alpha pathway (WP2878) and Thyroid stimulating hormone signaling pathway (WP2032) related to diabetes mellitus. Similarly, the moderate susceptible gene group were associated with pathways like Factors and pathways affecting insulin-like growth factor (IGF1)-Akt signaling (WP3850), Insulin signaling (WP481), Leptin-insulin signaling overlap (WP3935), Prolactin signaling pathway (WP2037), Roles of ceramides in the development of insulin resistance (WP5181), Thyroid stimulating hormone (TSH) signaling pathway (WP2032) and TNF-alpha signaling pathway (WP231) with significant impact on diabetes mellitus. Only three pathways like Leptin signaling pathway (WP2034), Prolactin signaling pathway (WP2037) and TNF-alpha signaling pathway (WP231) were found to be associated with the genes of less susceptible groups which have some associations with diabetes mellitus.

### 3.5 Protein centric annotations and selection of potential mutations in the candidate genes

The protein-centric information obtained from the UniProtKB database is used for annotation of genes described in high, moderate and less susceptible groups. The protein name, number of variants, number of predicted structures by different structure determination methods and protein family-related information were selected for the purpose (Supplementary file-2). All 77 candidate gene protein-centric annotations were thoroughly observed for their number of natural variants, structure predictions by different methods and familial information. About eight genes (ALB, ARRB2, CAV1, CAV2, CDH5, BNIP3L, CASP10 and FOXL2) were found to have no structural information, thus selected for computational modelling by different modelling protocols. Again the analysis of familial information revealed that about twenty-one genes (ACACB, ADARB1, AGER, AGRP, APAF1, APBB1, ARNTL, BARD1, BCL6, BID, BRCA1, CD4, CD44, CD47, CDH13, CREBBP, DAB2, DAG1, DFFA, JAG1 and KRIT1) were deprived of their familial information, thus selected for molecular modelling. A total of twenty-nine candidate genes were selected for computational model-based annotations for which their disease-related natural variants were selected in the lane of their potential mutations as reported by the PredictSNP algorithm.

### 3.6 Structural charactrization of candidate genes

Out of 77 candidate genes defined in three major groups about forty eight gene’s corresponding protein structural charactrization was done by the familial information obtained during protein centric annotations. Rest twenty nine gene’s corresponding protein structural annotation was done by the computational models designed by different methods.

#### 3.6.1 Familial information based

All forty-eight candidate genes were defined by thirty-two different families of information with peculiar structural and functional characters harboured by those group

##### ABC transporter superfamily, ABCA family

Only one candidate gene ABCA1 belong to this family. This family seems to be the largest known superfamily of proteins and members of this superfamily are involved in the hydrolysis of ATP and translocation of a broad spectrum of molecules across the cell membrane. About twelve members (ABCA1-ABCA13) of human ABC transporters have been reported to date which are full-sized proteins with a length of 1543 to 5058 amino acids.

##### Actin family

Two candidate genes ACTA1 and ACTG1 belong to this family. Members of this family are involved in wide range of cellular functions which include cell division, cellular migration, chromation remodelling and regulation of trancription and cell shape. Out of the mojor isoforms (ACTA1, ACTC1, ACTA2, ACTB, ACTG1) reported in mammals the protein encoded by ACTA1 expressed in skeletal tissue system and the protein encoded by ACTG1 is ubiquitously expressed in cytoplasm.

##### Alpha-actinin family

Two candidate genes ACTN2 and ACTN4 belong to this family which is a part of spectrin superfamily and charactrized by four known loci encoding four alpha-actins. About eight domains are present in members of alpha-actin family from which the actin binding domin seems to be highly conserved.

##### Apolipoprotein A1/A4/E family

only one candidate gene APOE belongs to this family. Apolipoproteins (apoA, apoC and apoE) function in lipid transport as structural components of lipoprotein particles, cofactors for enzymes and ligands for cell-surface receptors.apoE is a blood plasma protein that mediates the transport and uptake of cholesterol and lipid by way of its high-affinity interaction with different cellular receptors, including the low-density lipoprotein (LDL) receptor. These proteins contain several 22 residue repeats which form a pair of alpha helices.

##### Arrestin family

Two candidate genes named ARRB1 and ARRB2 belong to this small family of proteins. Arrestins are important for regulating the activity of G protein-coupled receptors(GPCRs) in the visual rhodopsin system and in the β-adrenergic system. The length of the arrestin family is about 404 amino acids. Both ARRB1 and ARRB2 are expressed at high levels in the central nervous system and may play a role in the regulation of synaptic receptors. Arrestin beta 2 was isolated from the thyroid gland, and thus it may also be involved in hormone-specific desensitization of TSH receptors.

##### BCL-2 family

Two candidate genes named BAD and BAX belong to this family.BCL-2 family proteins are the regulators of apoptosis, but also have other functions like neuronal activity, autophagy, calcium handling, mitochondrial dynamics and energetics, and other processes of normal healthy cells. BAD and BAX both are involved in the intrinsic mitochondrial apoptotic pathway and expressed in the transverse colon and other tissues. The BCL-2 family is divided into three groups based on their primary function (1) anti-apoptotic proteins (BCL-2, BCL-XL, BCL-W, MCL-1, BFL-1/A1), (2) pro-apoptotic pore-formers (BAX, BAK, BOK) and (3) pro-apoptotic BH3-only proteins (BAD, BID, BIK, BIM, BMF, HRK, NOXA, PUMA, etc.). Range in size from ∼100 to 200 amino acids.

##### Beta-catenin family

Only one candidate gene CTNNB1 comes under this family.β-catenin is an integral structural component of cadherin-based adherens junctions, and the key nuclear effector of canonical Wnt signaling in the nucleus.CTNNB1 is expressed in several hair follicle cell types: basal and peripheral matrix cells, and cells of the outer and inner root sheaths. Expressed in colon. Present in cortical neurons (at protein level). Expressed in breast cancer tissues (at protein level). 781 amino acid length has been found for the Beta-catenin family.

##### BZIP family

Three candidate genes ATF4, EBPA and CREB1 belong to this family. The basic leucine zipper (bZIP) gene family is known as transcription factor families. The Basic Leucine Zipper Domain is found in many DNA-binding eukaryotic proteins. bZIP transcription factors are effectors downstream of mitogenic stimulation, stress responses (oxidative, ER, heat), and cytokine stimulation. Additionally, the bZIP family of transcription factors affects several developmental processes including dendritic cell development, myeloid differentiation, and brain and ocular development. Based on protein homology and DNA-binding specificity, there are seven bZIP protein families, Jun, Fos, ATF, CREB, C/EBP, Maf, and are characterized by a highly conserved bZIP domain which is 60–80 amino acids.

##### Calcium channel alpha-1 subunit family

Only one candidate gene CACNA1I gene belongs to this family. This family is involved in a variety of calcium-dependent processes, including muscle contraction, hormone or neurotransmitter release, gene expression, cell motility, cell division, and cell death. The protein encoded by CACNA1I is highly expressed in brain tissue. The length of the protein is 2223 amino acids.

##### Calreticulin family

Only one candidate gene CALR gene belongs to this family. This family is involved in a Calcium-binding chaperone that promotes folding, oligomeric assembly, and quality control in the endoplasmic reticulum (ER) via the calreticulin/calnexin cycle. The protein encoded by CALR is extensively expressed in all eukaryotic organisms. The length of the protein is 417 amino acids.

##### Caveolin family

Three candidate genes CAV1,CAV2, and CAV3 belong to this family. This family is involved in the costimulatory signal essential for the T-cell receptor (TCR)-mediated T-cell activation. The caveolin family of proteins is encoded by three genes and consists of six known caveolin subtypes: caveolin-1α, caveolin-1β, caveolin-2α, caveolin-2β, caveolin-2γ, and caveolin-3. Of the six known caveolins, caveolin-3 is specifically expressed in muscle tissue, including the heart. The length of the proteins CAV1, CAV2 and CAV3 are 147,162 and 151 amino acids.

##### Clusterin family

Only one candidate gene CLUH gene belongs to this family. This family is involved in several basic biological events such as cell death, tumor progression, and neurodegenerative disorders. The protein encoded by the CLUH gene is a secreted chaperone that can under some stress conditions also be found in the cell cytosol. The length of the protein is 1309 amino acids.

##### CRK family

Only one candidate gene CRKL gene belongs to this family. This family is involved in the transduction of intracellular signals. The protein encoded by CRKL is highly expressed in secondary oocytes and other tissue. The length of the protein is 303 amino acids.

##### DAXX family

only one candidate gene death-associated protein-6(DAXX) belongs to this family.DAXX modulates transcription through binding to transcription factors, epigenetic modifiers, and chromatin remodelers. DAXX’s localization in the PML nuclear bodies also plays role in transcriptional regulation. The size of amino acid residues in the DAXX family is 740.

##### DEAD box helicase family, DDX3/DED1 subfamily

DDX3X and DHX9 are the two candidate genes belonging to this family. Dead-box proteins are ATP-dependent RNA-binding proteins that remodel RNA structures and RNA-protein complexes, stably clamp RNA, and promote fluidity within RNA granules.DDX3X Suppresses the Susceptibility of Hindbrain Lineages to Medulloblastoma. DHX9 is a multi-domain, multi-functional protein, with regulatory roles in DNA replication, transcription, translation, RNA processing, and transport, microRNA processing, and maintenance of genomic stability.

##### Fibrillar collagen family

only one candidate gene COL1A1 belongs to this family. Fibril-associated collagen with interrupted triple helix (FACIT) collagens (fibril-associated collagens with interrupted triple-helices), include types IX, XII, XIV, XIX, XX, and XXI. Many FACIT collagens associate with larger collagen fibers and act as molecular bridges stabilizing the organization of the extracellular matrices. The COL1A1 gene provides instructions for making part of a large molecule called type I collagen. Collagens are a family of proteins that strengthen and support many tissues in the body, including cartilage, bone, tendon, skin, and the white part of the eye. The prototype collagen (type I) has an uninterrupted Gly-X-Y repeat sequence that is almost 1000 amino acid residues in length.

##### G-protein coupled receptor 1 family

ADORA1 and APLNR are the two candidate gene belonging to G-protein coupled receptor 1 family. Family 1 G protein-coupled receptors (GPCRs) are activated by a large number of ligands including photons, odorants, neurotransmitters, and hormones, and are involved in a wide variety of central and peripheral functions. GPCRs 1 family plays a major role in the regulation of neuronal activity and behavior. *ADORA1* (Adenosine A1 Receptor) is a Protein Coding gene with adenosine *as an endogenous ligand*.ADORA1 regulates tumor PD-L1 expression. APLNR is related to the angiotensin receptor but is actually an apelin receptor that inhibits adenylate cyclase activity and plays a counter-regulatory role against the pressure action of angiotensin II by exerting a hypertensive effect. The amino acid size of this family varies between 290 and 951 amino acid residues.

##### GADD45 family

GADD45 is the only candidate gene coming under this family. GADD45 family which includes GADD45A, GADD45B, and GADD45G implicated as stress sensors that modulate the response of mammalian cells to genotoxic and physiological stress and modulate tumor formation. The length of amino acid residues of this family ranges from 150 to 165 amino acids.

##### Glucagon family

only one candidate gene ADCYAP1 belongs to this family. The glucagon family peptides include glucagon, the glucagon-like peptides (GLP-1 and GLP-2), and GIP, which is a subfamily within the secretin family and shares a highly related molecular structure. The peptides are expressed in endocrine cells of the pancreas and gastrointestinal epithelium, and in specialized neurons in the brain, which play a major role in the regulation of nutrient homeostasis and energy metabolism through their distinct specific receptors. ADCYAP1 Binding to its receptor activates G proteins and stimulates adenylate cyclase in pituitary cells. Promotes neuron projection development through the RAPGEF2/Rap1/B-Raf/ERK pathway. In chromaffin cells, induces a long-lasting increase in intracellular calcium concentrations and neuroendocrine secretion. Involved in the control of glucose homeostasis, induces insulin secretion by pancreatic beta cells. GLP-1 and its family peptides are 30 to 40 amino acids in length.

##### IAP family

Two candidate genes BIRC2 and BIRC3 belong to the IAP family. Inhibitor of apoptosis (IAP) proteins interface with, and regulate a large number of, cell signaling pathways.IAP family proteins are characterized by a novel domain of ∼70 amino acids.BIRC2 and BIRC3 play various roles in anti-apoptotic, tumor necrosis factor (TNF) mediated, and canonical/non-canonical NFkB signaling.

##### Lipoxygenase family

Only one candidate gene ALOX5 belongs to the Lipoxygenase family. Lipoxygenases form a family of lipid peroxidizing enzymes, which oxygenate free and esterified polyenoic fatty acids to the corresponding hydroperoxy derivatives. Catalyzes the oxygenation of arachidonate to 5-hydroperoxyeicosateraenote followed by the dehydration to 5,-epoxyeicosatetraenote, the first two steps in the biosynthesis of leukotrienes the first of which are potent mediators of inflammation. lipoxygenase family consisting of 662–711 amino acids.

##### MAM33 family

Only one candidate gene C1QBP gene belongs to this family. This family is involved in inflammation and infection processes, ribosome biogenesis, protein synthesis in mitochondria, regulation of apoptosis, transcriptional regulation, and pre-mRNA splicing. The protein encoded by C1QBP is highly expressed in the cell surface of peripheral blood cells at the protein level and Surface expression is reported for macrophages and monocyte-derived dendritic cells. The length of the protein is 282 amino acids.

##### MIP/aquaporin family

Two candidate genes AQP and AQP12B belong to this family. Members of the family are involved in forming a water-specific channel that provides the plasma membranes of red cells and kidney proximal tubules with high permeability to water, thereby permitting water to move in the direction of an osmotic gradient. Out of the major isoforms AQP1, AQP4, AQP7, and AQP12B reported in mammals the protein encoded by AQP1 and expressed in adipose tissue, liver, and endocrine pancreas. The length of the protein is 269 to 323 amino acids.

##### NIP3 family

Only one candidate gene NECAME gene belongs to this family. This family is involved in Aspartic protease which cleaves several human serum proteins including hemoglobin, fibrinogen, and albumin. Appears to cleave preferentially between P1 (Ala, Leu, Val, Phe, and Gly) and P1’ (Ala and Leu) residues. The protein encoded by NECAME is highly expressed in the intestine, amphidal glands, and excretory glands. The length of the protein is 425 amino acids.

##### Peptidase C14A family

Only one candidate gene CASP2 gene belongs to this family. This family is involved in mediating cell death (apoptosis). The protein encoded by CASP2 is highly expressed in neurodegenerative disorders including Alzheimer’s disease, Huntington’s disease, and temporal lobe epilepsy. The length of the protein is 452 amino acids.

##### PI3/PI4-kinase family, ATM subfamily

Only one candidate genes ATM gene belong to this family. This family is involved in regulating DNA damage response mechanism, signal transduction, and cell cycle control, and function as a tumor suppressor, The protein encoded by ATM is highly expressed in lymphoid tissue. The length of the protein is 3056 amino acids.

##### Protein kinase superfamily, AGC Ser/Thr protein kinase family

Only one candidate gene PRKCI gene belongs to this family. This family is involved in Protein phosphorylation, which plays a key role in most cellular activities, and is a reversible process mediated by protein kinases and phosphoprotein phosphatases. The protein encoded by PRKCI is highly expressed in blood and adipose tissue. The length of the protein is 596 amino acids.

##### TRAFAC class dynamin-like GTPase superfamily

Two candidate genes DNM1L and IFGGE gene belong to this family. This family is involved in the mitochondrial and peroxisomal division. Out of the major isoforms (DNM1L, DNM1, DNM2, DNM3, ATL, IFGGE) reported in mammals. The protein encoded by DNM1L is expressed in Granular cytoplasmic and membranous expression in several tissues and the protein encoded by IFGGE is ubiquitously expressed in the cytoplasm.

##### Tumor necrosis factor family

Only one candidate gene FASLG belongs to this family which is described as a large family of proteins. About 17 members of this family are reported to date which have significant roles in inflammation, immunity and a broad range of diverse biological functions.

##### Vacuolar ATPase subunit S1 family

Only one candidate gene ATP6AP1 belongs to this family. This family comprises highly conserved evolutionary significant enzymes with diverse functions in eukaryotes. The major functions of these member proteins include copulation of ATP hydrolysis energy to the proton transport system across intracellular and plasma membranes of eukaryotes.

#### 3.6.1 Computational model based

The computational models predicted for twenty nine candidate genes were checked for their structural stability by generating Ramachandran plots. Almost all the models were observed to have 90% amino acid residues in the allowed region and in some cases this criterion is fulfilled by considering the residues occupied in the allowed and additional allowed regions. The analysis of all the predicted models is summarized in Supplementary file-3.

The structural characterization of corresponding candidate genes was done by analysing their computational models and fetching of domain information if any. The models generated by homology modelling were characterized by the domain information defined by their templates during database similarity search. The models generated by the I-TASSER threading server were characterized by the structural information only.

## 4. Conclusion

In the lane of identification, validation and structural characterization of potential candidate genes for diabetes mellitus we have integrated validated information from various databases like HIPPIE, DisGeNET, GEO, STRINGS, UniProtKB and PDB using several computational algorithms of systems biology. The result was quite generous with the identification of about 89 candidate genes of which only 77 genes were appropriate to our validation protocol as they were in the close proximity regions of diabetes mellitus-related genes. Again the structural characterization of these candidate gene proteins revealed important information that can be used in understanding the aetiology of this complex disorder and the structure-based design of new chemical entities (NCEs) for its treatment.

## 5. Acknowledgements

The authors are thankful to the Vice-Chancellor of Utkal University, Head of Department of University Department of Pharmaceutical Sciences and Faculties of Bioprudence Research Innovations LLP for providing essential facilities. The award of ICMR-SRF Fellowship for Dhananjay Kumar Tanty and UGC Fellowship for Prachi Rani Sahu are duly acknowledged.

## Notes

### Competing Interest Statement

The authors have declared no competing interest.

